# Molecular dynamics simulation of proton-transfer coupled rotations in ATP synthase F_O_ motor

**DOI:** 10.1101/618504

**Authors:** Shintaroh Kubo, Toru Niina, Shoji Takada

**Affiliations:** Department of Biophysics, Graduate School of Science, Kyoto University, Kyoto 606-8502, Japan

## Abstract

The F_O_ motor in F_O_F_1_ ATP synthase rotates its rotor driven by the proton motive force. While earlier studies elucidated basic mechanisms therein, recent advances in high-resolution cryo-electron microscopy enabled to investigate proton-transfer coupled F_O_ rotary dynamics at structural details. Here, developing a hybrid Monte Carlo/molecular dynamics simulation method, we studied reversible dynamics of a yeast mitochondrial F_O_. We obtained the 36°-stepwise rotations of F_O_ per one proton transfer in the ATP synthesis mode and the proton pumping in the ATP hydrolysis mode. In both modes, the most prominent path alternatively sampled states with two and three deprotonated glutamates in c-ring, by which the c-ring rotates one step. The free energy transduction efficiency in the model F_O_ motor reaches ~ 90% in optimal conditions. Moreover, mutations in key glutamate and a highly conserved arginine increased proton leakage and markedly decreased the coupling, in harmony with previous experiments.

## Introduction

F_O_F_1_ ATP synthase is a ubiquitous protein complex that plays the central role in the energy generation of living cells (*1*, *2*). The enzyme complex is located at the inner membrane of mitochondria, the thylakoid membrane of chloroplast, and the bacterial membrane, working for most of ATP production in cells. Dysfunction in the ATP synthase causes severe diseases such as neuropathy, maternally inherited leigh syndrome, ataxia and retinitis pigmentosa syndrome (*3*–*5*). F_O_F_1_ ATP synthase is composed of two regions, F_O_ embedded in the membrane and F_1_ protruded to the matrix solution (Fig.1A); each of the regions works as a reversible rotary motor (*2*, *6*). Driven by the proton motive force, the proton flux through the F_O_ motor from the inter membrane space (IMS) to the matrix generates a torque in the rotor. The torque is transmitted to the central rotor in F_1_ where the rotary motion is used to synthesis ATP (*7*, *8*). Conversely, *in vitro*, and *in vivo* for bacteria, the F_1_ motor can hydrolyze ATP to generate a torque in the inverse direction and drives F_O_ to pump up protons.

**Fig. 1.**
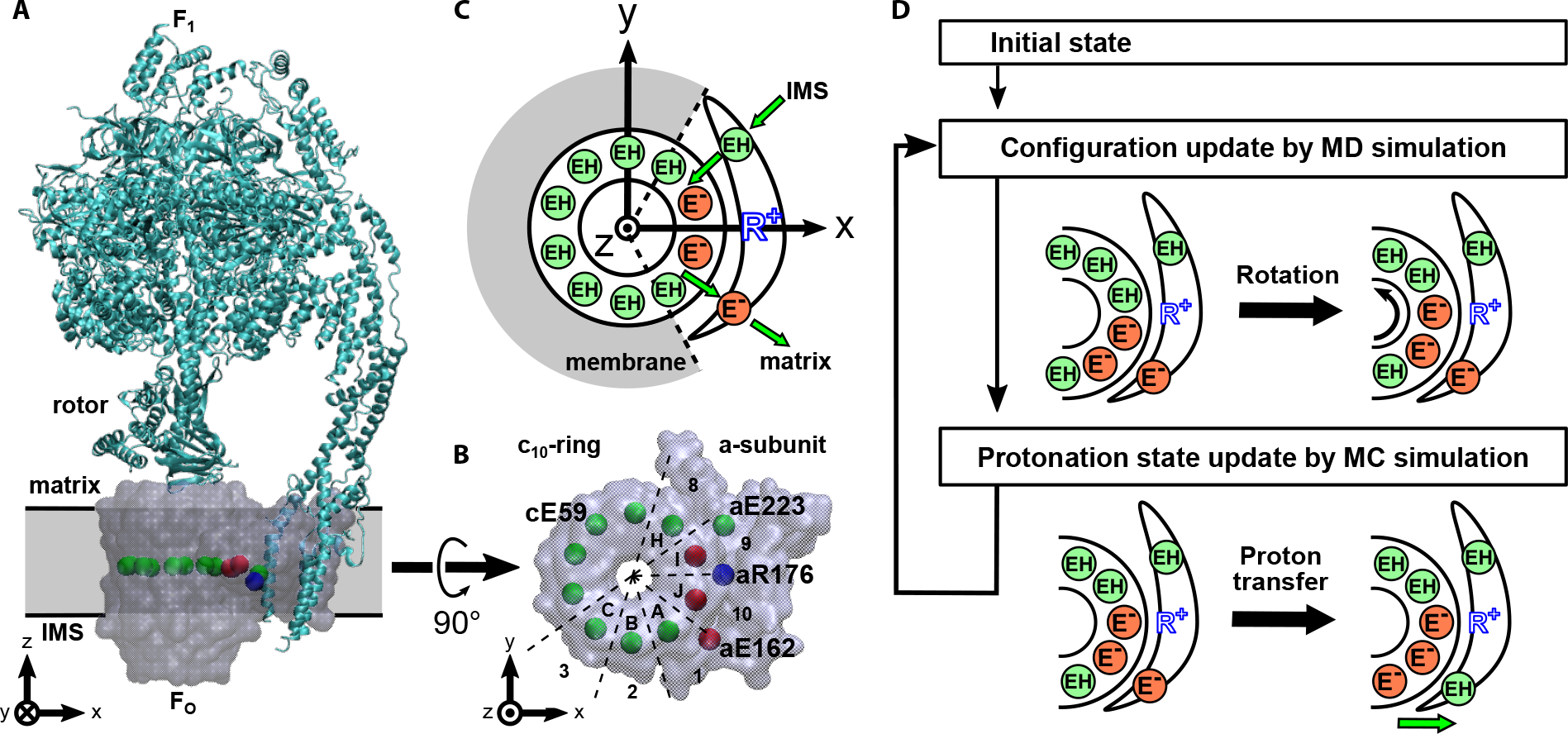
Simulation setup. **(A)** Side view of the yeast F_O_F_1_ ATP synthase whole complex (PDB ID: 6CP6). The ac_10_ part of F_O_ region is depicted as the transparent surface, while the F_1_ region and the b-subunit are in the ribbon diagram. The rectangular gray region represents the membrane. The beads inside F_O_ region represent cE59, aE223, aE162, and aR176 (blue). Putatively protonated and deprotonated glutamates are drawn in green and red, respectively. **(B)** Top view of our simulation system, ac_10_ in the F_O_. We assign alphabets A to J to identify each c-subunit in c_10_-ring in clockwise order starting from the nearest chain from aE162, which connects to the matrix region. aE223 is connected to the IMS side. **(C)** Schematic picture of our model. EH and E^−^ represent protonated and deprotonated glutamates, respectively. Membrane drawn in gray is modeled implicitly. Protons can transfer between cE59 and glutamates in a-subunit, aE223 and aE162. Also, aE223 and aE162 exchanges their protons with the IMS and the matrix aqueous environment, respectively. Arrows in green indicate the net proton flow in the ATP synthesis mode. We set the rotational axis of the c_10_-ring as the z-axis and the position of aR176 on the x-axis. **(D)** Our simulation protocol. We performed short MD and MC simulations in turn. MD simulation updates the protein structure and MC simulation updates the protonation states of 12 glutamates.

The motor function of F_O_ arises from a complex of c-ring and a-subunit (Fig.1B). The c-ring is made up of a varying number of c-subunits from 8 to 17 depending on the species (*6*, *9*–*14*). Here, we focus on yeast mitochondrial F_O_ having 10 c-subunits in the c-ring, which we call c_10_-ring (*10*, *15*). The c-subunit has a key glutamate (cE59, throughout, the residue number is for yeast F_O_F_1_) nearly at the middle point in the membrane, which serves as the carrier of protons: A protonation in the cE59 allows c_10_-ring to hold a proton, whereas it is deprotonated upon a proton release. It is experimentally confirmed that the bacterial ATP synthase activity drastically decreases when the corresponding key residue is mutated into the amino acids that cannot carry a proton (*16*, *17*). The a-subunit is considered to be a mediator between the aqueous environment and c-ring: The a-subunit has two separate half-channels; one connecting c-ring to the IMS side, and the other connecting c-ring to the matrix side. The recent cryo-electron microscopy (cryo-EM) studies (*18*, *19*) revealed the structure of F_O_F_1_ ATP synthase at nearly the atomic resolution and found two long tilted parallel *α*-helices in the a-subunit at the interface with c_10_-ring. On each side of the helices, two half-channels are located. These half-channels in the a-subunit have highly conserved glutamate residues, aE223 and aE162, which are considered as proton relaying sites (*20*). Additionally, the arginine residue aR176 at the middle of the long parallel helices plays also an important role to separate two half channels by preventing protons from leaking. It is experimentally confirmed that the mutation in this key arginine leads to proton leakage and decreases both the ATP synthesis and the proton pump activities for the thermophilic F_O_F_1_ (*21*). Additionally, because aR176 locates close to cE59 in c_10_-ring, it is speculated that the attractive interaction between aR176 and deprotonated cE59 also contributes to increase the efficiency of the F_O_ rotation (*22*). However, it is impossible to observe the proton transfers through F_O_ directly.

Based on these knowledge, numerous models for the working mechanisms in F_O_ have been proposed. Vik and Antonio first proposed the widely-accepted model that two half-channels in a-subunit shuttle protons with the proton carrier acid residues (cE59 for yeast) in the middle of c-ring, by which c-ring rotates relative to a-subunit (*22*). Subsequently, this model was extended to a mathematical and quantitative model by Oster and colleagues (*23*). In these models, in addition to basic proton flows, switches in electrostatic attractions between protonated and deprotonated acidic residues with a conserved arginine provide key controls in functions. These together successfully explains many experimental data. Many subsequent theoretical works are more or less based on these models (*24*). However, since these models do not consider the atomic structure explicitly and, thus unavoidably, how proton transfer is coupled with rotation was not pursued in structural detail.

Here, we address the proton-transfer coupled F_O_ rotary motions at high resolution, developing a generic computational framework to simulate the protein structure dynamics and proton transfers simultaneously. We employ Metropolis Monte-Carlo (MC) method to simulate proton transfers and combine it to molecular dynamics (MD) simulations. We performed this hybrid MC/MD simulation, in which MD stages for protein dynamics and MC stages for proton transfers are alternatively conducted (Fig.1D). In MD simulations, for computational ease, we used a well-established coarse-grained model in which each amino acid is represented by one bead (*25*, *26*). We modeled the protonation state by the value of charges on each amino acid. We applied an implicit membrane potential that gives rise a free energy cost when deprotonated glutamates face to lipid membrane region (Fig.S1A) but not the interface with the a-subunit (Fig.1C). Our simulations, for the first time to our knowledge, successfully reproduced the proton-transfer coupled F_O_ rotation (i.e., the ATP synthesis mode) and, conversely, the torque-driven proton pumping (i.e., the ATP hydrolysis mode), with explicit molecular structures. We identified several prominent pathways of how proton-transfer couples the c-ring rotation, as well as of proton leakage. By mutating cE59 and aR176, we clarified the behavior of the mutants that lack ATP synthesis and/or proton pump activity. Additionally, we investigated behaviors of F_O_ in various different physical parameters, such as external torque, pH, membrane potential, and the intrinsic pKa of glutamate residues.

## Results

### F_O_ rotation driven by the proton-transfer across the membrane

We begin with simulations of rotary motions of F_O_ motor driven by a proton-motive force, which corresponds to the ATP synthesis mode. Without external torque applied to the c_10_-ring, we performed proton-transfer coupled MD simulations of the F_O_ motor ac_10_ with the pH difference between the IMS and matrix sides being 1 and the membrane potential ΔΨ = 150mV. Using the hybrid MC/MD method (see Materials and Methods), we carried out 10 independent simulations with stochastic forces. Fig.2A shows cumulative rotational angles of the c_10_-ring in each trajectory together with the number of protons that are transferred from the IMS to the matrix throughout (a representative trajectory in red corresponds to Movie S1). Clearly, we see that F_O_ rotates stochastically and unidirectionally, and that protons were transferred as F_O_ rotates. Note that here we counted the number of protons that transferred all the way from the IMS to the matrix and thus the protons that were bound in F_O_ in the initial state were not included.

**Fig. 2.**
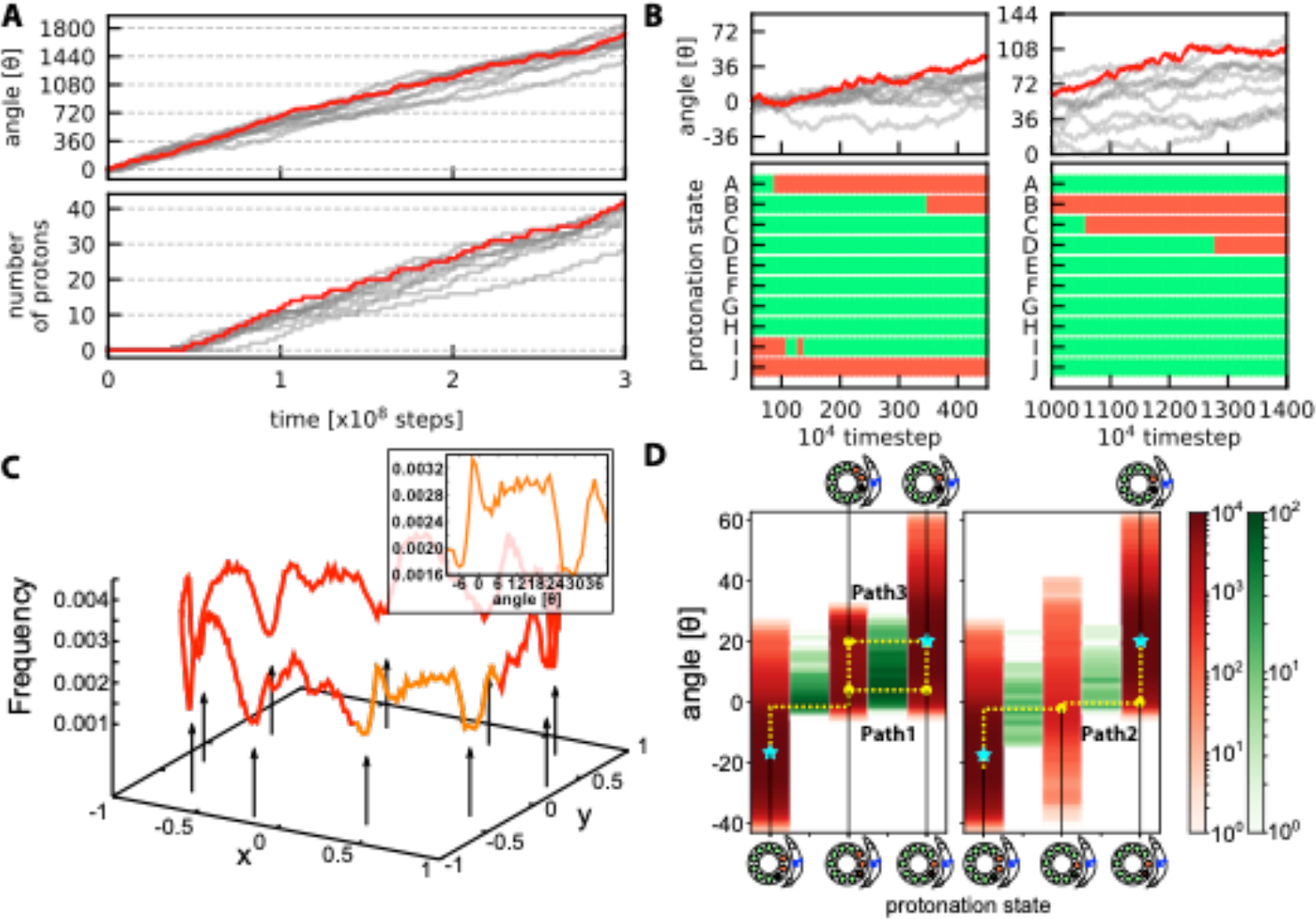
Simulations driven by a proton-motive force for the ATP synthesis mode. **(A)** Time courses of the cumulative rotation angle of c_A_E59 (top) and the number of protons that across all the way from IMS side to matrix side (bottom) for 10 trajectories. The positive rotation angle means the counterclockwise rotation corresponding to ATP synthesis mode. A representative one is in red, whereas the other 9 in gray. **(B)** A closeup view of the rotation angle (top) together with a time course of the protonation states of 10 cE59’s in the representative case (the red trajectory in the top) (bottom). The green (red) mean protonated (deprotonated) states. **(C)** Probability distribution of the angle of c_A_E59 in the representative trajectory (the red one in A and B). Arrows indicate low probability angles. The inset shows the closeup view of the orange region. inset; xy plot for one rotation state. **(D)** Protonation state dependent distribution of the cumulative rotation angle (red) and the count of transition events from the left to the right side (green) for different pathways. (Left) Paths 1 and 3. From the left, each red lane represents, DEP(c_I_, c_J_), DEP(c_I_, c_J_, c_A_), and DEP(c_J_, c_A_). (Right) Path 2. From the left, each red lane represents, DEP(c_I_, c_J_), DEP(c_J_), and DEP(c_J_, c_A_). Cyan stars represent stable states. Yellow curves illustrate three pathways.

The averaged net number of protons that passed from the IMS to matrix sides, the proton flux, per c_10_-ring revolution was ~10, which clearly suggests the stoichiometric tight coupling between the proton transfer and the rotation (Table S1).

### F_O_ rotates stepwise by 36° coupled with a proton transfer from IMS to matrix

Since the architecture of the c_10_-ring has a 10-fold symmetry, it is thought that F_O_ rotates in 36° steps, and the existence of such steps has been confirmed for *E. coli* F_O_F_1_ (*27*). In Fig.2B, we find the time when the c_10_-ring rotary angle comes over every 36° and the time at which the protonation state changes are synchronized although the rotation steps are not very clear in our simulations. Then, we plotted the population of c_A_-chain angle around the rotational axis from 10 trajectories (Fig.2C), finding that there are 10 low-population gaps in symmetric locations that separate 10 states (the black allows in Fig.2C). This is in a good agreement with the 10 symmetric states observed in the experiment (*27*). Notably, the population of c_A_ angle has relatively broad distribution separated by narrow unstable gaps. The inset of Fig.2C shows the orange colored plot in Fig.2C with x-axis being rotational angle of c_A_-chain. From this figure, we find the angles at −6° and 30° correspond to low-population angles (the angle zero corresponds to the initial state). Thus, the c_10_-ring does rotate stepwise by ~ 36°. The 36° step is mainly caused by the low-population bottleneck angles.

Additionally, in the experiment (*27*), 72° steps were observed. One possible explanation about why they observed such a long step is the low time resolution compared to the F_O_ rotation. Our result suggests that the thermal fluctuation can occasionally lead F_O_ to rotate faster and the successive two transitions can be interpreted as one, 72° step.

### Pathways in the proton-motive force driven F_O_ rotation

Next, we analyzed detailed pathways of proton-transfer coupled rotary motions in the above simulations. Classifying the pathways, we obtained a few major routes in the ATP synthesis mode (Fig.2D). Taking advantage of the 10-fold symmetry, here we focus on one step of proton-transfer coupled rotation by 36°. Transitions from the previous step occur around θ~ − 18°, relative to the angle in the cryo-EM. Given that, the transition pathway starts with the state {DEP(c_I_, c_J_), θ~ − 18°}, in which only c_I_ and c_J_ in c-ring are deprotonated. In the most prominent path (Fig.2D left), firstly, upon a counterclockwise rotary fluctuation up to θ~ − 2° from θ~ − 18°, the protonated c_A_ gives a proton to the matrix site aE162 and becomes deprotonated, {DEP(c_I_, c_J_, c_A_), θ~ − 2°}. Subsequently, it rotates up to θ~3°, and then, the deprotonated c_I_ receives a proton from the IMS site aE223 and becomes protonated {DEP(c_J_, c_A_), θ~3°}. Then, the c_10_-ring further rotates counterclockwise, resulting in {DEP(c_J_, c_A_), θ~18°}. This final state is equivalent to the initial state with a shift in the chain identify. We call this pathway as Path 1 hereafter.

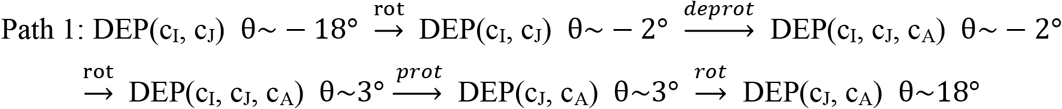

Additionally, we found sub-dominant pathways, Path 2 (Fig.2D right) and Path 3 (Fig.2D left). In Path 2, the protonation of c_I_ occurs before the deprotonation of c_A_^1^.

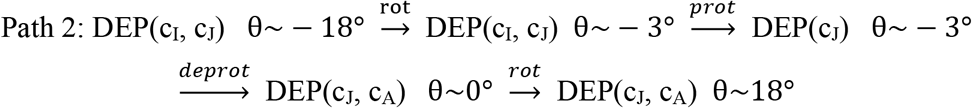

The Path 3 is a small variant of Path 1; the second half of F_O_ rotation occurs before the protonation of the c_J_.

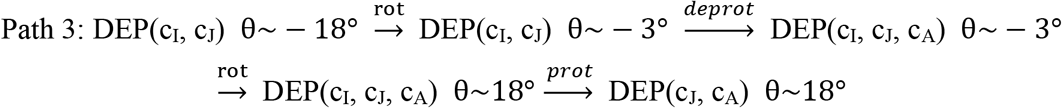

### The external torque can pump the proton across the membrane

F_O_F_1_ ATP synthase can work as a proton pump driven by the ATP hydrolysis. ATP hydrolysis in F_1_ leads to the clockwise rotation of the central stalk, which transmits the torque to F_O_ c_10_-ring. The rotation in F_O_ c_10_-ring results in pumping protons from the matrix side to the IMS side (Note that, for yeast mitochondrial F_O_F_1_, the proton pump occurs only *in vitro*, but we use the terms, the matrix side and the IMS side for simplicity). Here, we investigated the proton-pump function of F_O_ driven by an external torque applied to c_10_-ring.

In this study that contains only the F_O_ part, we applied an external torque to the residues on the top of c_10_-ring, cS38, by which we mimic the work by the ATP hydrolysis in F_1_. We turned off the proton-motive force by setting pH in both sides equal and the membrane potential ΔΨ = 0mV. Fig.3A shows 10 trajectories of rotation angles and the number of protons pumped from the matrix to the IMS environment under 86.2 pN · nm torque, which corresponds to the F_1_ torque observed in experiments (*28*) (the proton flux from the matrix to IMS sides is defined as the negative value)(a representative trajectory in red corresponds to the Movie S2). Clearly, protons are pumped up stochastically and unidirectionally across the membrane driven by mechanical forces. At this condition, the average number of pumped protons per revolution was ~10 (Table S1).

**Fig. 3.**
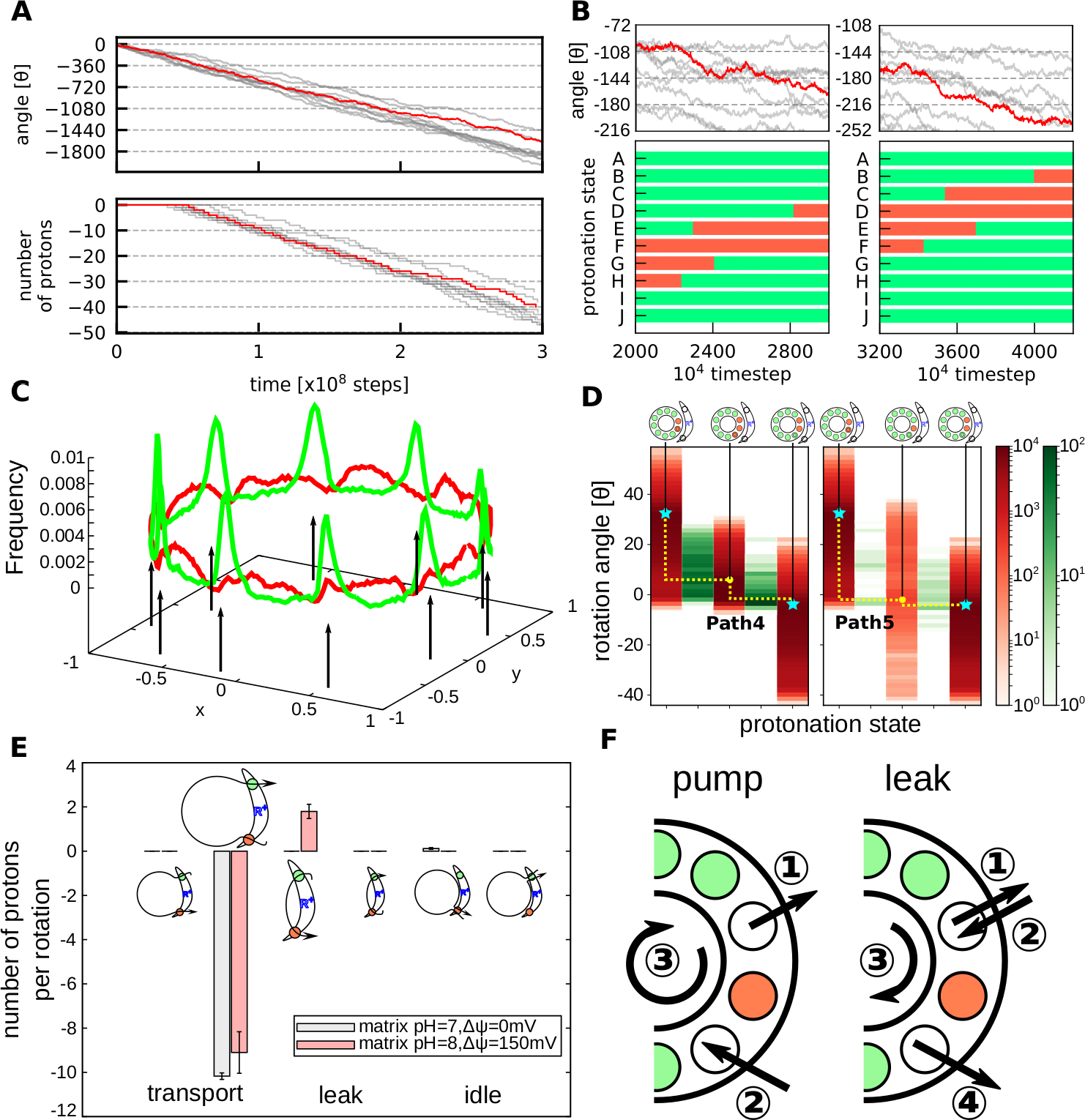
Simulations driven by an external torque for the ATP hydrolysis mode. **(A)** Time courses of the cumulative rotation angle of c_A_E59 (top) and the number of protons that across all the way from IMS side to matrix side (bottom) for 10 trajectories in the absence of the proton-motive force. The negative rotation angle means the clockwise rotation corresponding to proton pumping. A representative one is in red, whereas the other 9 in gray. **(B)** A closeup view of the rotation angle (top) together with a time course of the protonation states of 10 cE59’s in the representative case (the red trajectory in the top) (bottom). The green (red) mean protonated (deprotonated) states. **(C)** Probability distribution of the angle of c_A_E59 in the representative trajectory in green (the red one in A and B). Arrows indicate low probability angles. For comparison, the red curve is the result from ATP synthesis mode (Figure 2C). **(D)** Protonation state dependent distribution of the rotation angle (red) and counts of transition events from the left to the right sides (green) for different pathways. (Left) Path 4. From the left, each red lane represents, DEP(c_I_, c_J_), DEP(c_I_, c_J_, c_A_), and DEP(c_J_, c_A_). (Right) Path 5. From the left, each red lane represents, DEP(c_I_, c_J_), DEP(c_J_), and DEP(c_J_, c_A_). Cyan starts are stable states. Yellow curves illustrate three pathways. **(E)** Numbers of protons transported per 36° rotation. The positive values mean that the transport from IMS to matrix side and vice versa. “transport”; the protons contributes to the rotation of c_10_-ring. “leak”; the protons do not contribute to the rotation of c_10_-ring but transferred across the membrane, and “idle”; the protons do not contribute to the rotation and come back to the same side of the bulk. **(F)** Schematic view of how the protons leak without contributing to the rotation.

Compared to the ATP synthesis process shown in Fig.2B, there is a clearer stepwise motion in 36° (Fig.3BC). It is because the applied torque pushes the c_10_-ring to near the unstable gap found in the ATP synthesis mode, resulting in a sharp and high population there.

Next, we move to a tug-of-war situation: We applied the 86.2 pN · nm external torque to F_O_ with the standard proton-motive force; the pH difference being 1 and the membrane potential ΔΨ = 150 mV. At this condition, the external torque is expected to be stronger than the torque generated by the proton-motive force. Indeed, the simulation resulted in proton pumping coupled with the clockwise rotation of c-ring. Here, the average number of protons pumped per revolution was about 9 with some proton leakage (Fig.3E). The proton leakage occurs in the following way. Firstly, in the functional pumping process, proton escapes from c_10_-ring to the IMS side. Subsequently, a proton enters from matrix side into c_10_-ring and it rotates (Fig.3F left). Occasionally, however, after the proton escape from c_10_-ring to the IMS side, another proton comes back to c_10_-ring from the IMS side. This proton occasionally reaches to the matrix side, resulting in the leakage from IMS to matrix side (Fig.3F right). Compared with the functional proton pumping, it can be said that the leakage occurs if a proton enters into c_10_-ring from IMS side before a proton comes from matrix side and stays there until the ring rotates.

### Pathways in the proton pump motion

Fig.3D shows major pathways observed in the simulations of the ATP hydrolysis mode in the absence of the proton-motive force. For simplicity, we start from the state in which c_J_ and c_A_ are deprotonated {DEP(c_J_, c_A_), θ ~ 34°}. Notably, even though the protonation state DEP(c_J_, c_A_) has its intrinsic free-energy minimum at θ ~ 36°, the external torque skews the free energy surface so that the minimum shifts to θ ~ 0°. In this initial state (Fig.3D left), c_I_ frequently exchange its proton with the IMS site aE223. If c_A_ receives a proton from matrix side aE162 while c_I_ is protonated, the protonated c_A_ can immediately enter into the membrane, driven by the torque applied to c_10_-ring, and F_O_ rotates in the clockwise direction by ~ 36°. Then the protonation state returns to the state equivalent to the initial state with a shift in the chain identify. We call this as Path 4.

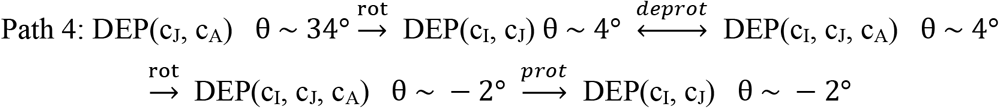

Also, a sub-dominant pathway Path 5 in which c_A_ is protonated before c_I_ is deprotonated was observed (Fig.3D center).

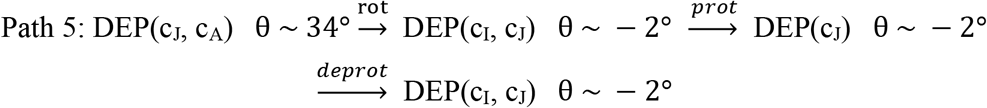

The Path 4 is more frequent than the Path 5 because it is easier to receive a proton from the IMS side than from the matrix side due to the difference in the local pH and the membrane potential.

Next, we investigate the pathways in the presence of the proton-motive force as well as the same external torque, i.e., the tug-of-war situation. Results in Fig.S2 shows that the proton pump activity is much slowed down (Fig.3E). As described above, we found some proton leak, as well. Since the proton exchange between c_10_-ring and the IMS site is fast, F_O_ occasionally have a proton in c_I_ when c_A_ is protonated. After the protonation at c_A_, F_O_ can quickly be rotated 36° in the clockwise direction by the applied torque, by which a proton escapes to the matrix side immediately. In this pathway Path 6, the proton does not go around c_10_-ring and causes proton leakage.

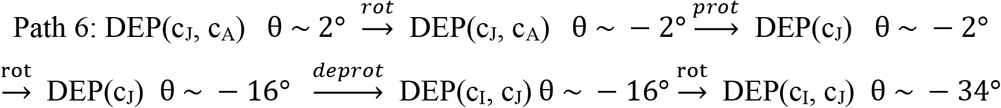

In summary, given the two dominant paths, the Path 1 and the Path 4, there is a tendency that deprotonation of the c_10_-ring occurs before protonation in both ATP synthesis and hydrolysis processes and subsequently F_O_ rotates either in counterclockwise (in the ATP synthesis mode) or clockwise direction (in the ATP hydrolysis mode).

### Mutations in F_O_

So far, we have described the behavior of the WT F_O_ motor. Now, we address effects of some mutations on critical residues; glutamic acids in c_10_-ring and the aR176. It has been experimentally confirmed that a bacterial F_O_F_1_ loses its proton pump function when the corresponding arginine is mutated to alanine(*21*). In addition, it has also been confirmed that the ATP synthesis activity of a bacterial F_O_F_1_ drastically decreases when the acidic residue equivalent to cE59A in one of c-subunits is mutated (*17*). In order to investigate the mechanism how these mutations lose the functionality, we carried out MC/MD simulations for these mutants.

### The aR176A mutant rotates with some protons leak in the ATP synthesis mode

First, we modeled the mutant in which aR176 is substituted with alanine (aR176A) and performed the hybrid MC/MD simulation with no external torque in the same way as above. Results show that, while the unidirectional rotation does occur, the rotation for aR176A was clearly slower than that of WT (Fig.4A). Also, the number of transported protons per revolution was about 10 (Fig.4C).

**Fig. 4.**
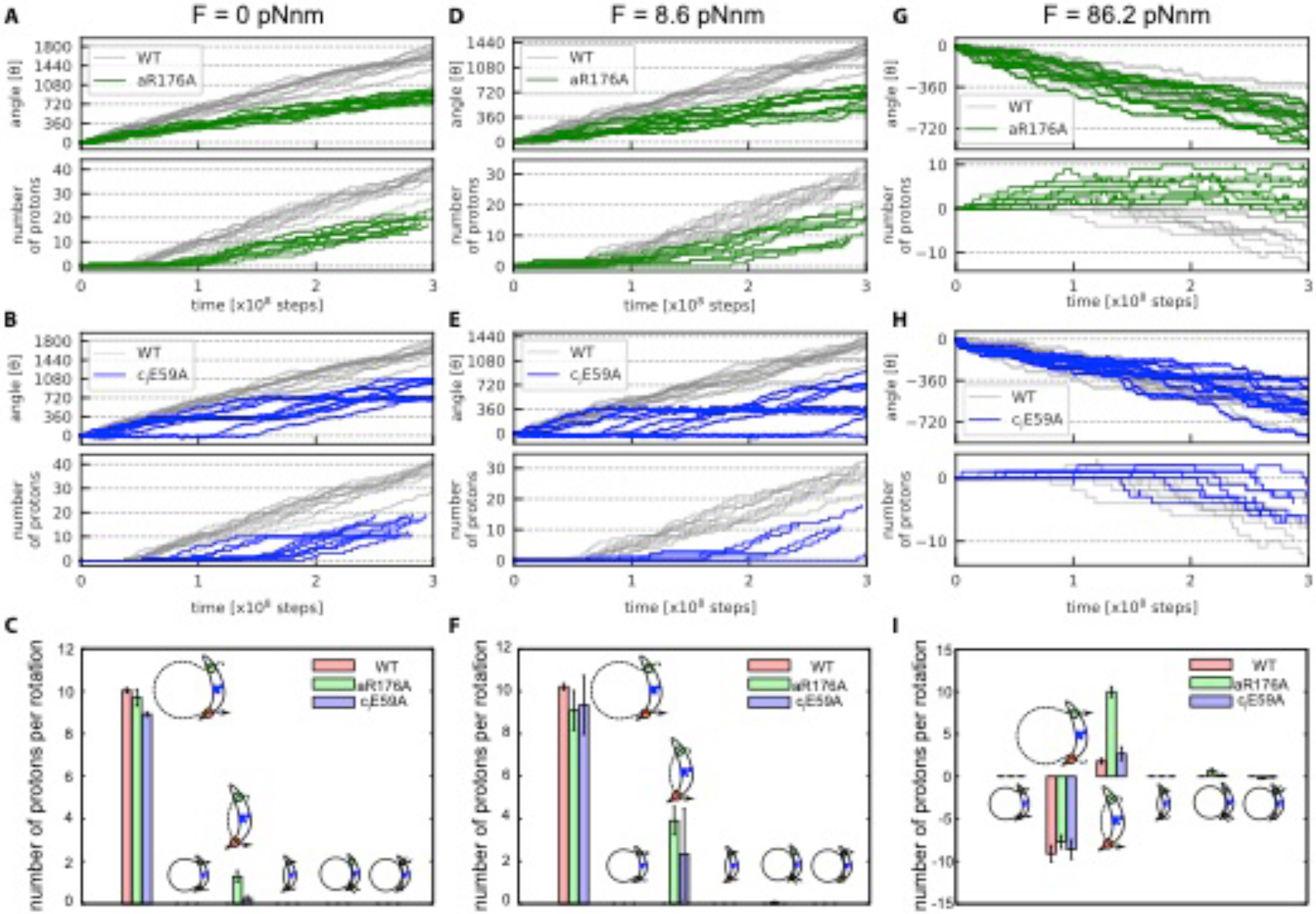
Simulations of mutant F_O_’s. Rotary motions, proton transfers and pathways are plotted for two mutant F_O_’s, aR176A (green curves in **A, D, G**) and c_J_E59A (blue curves, **B, E, H**), with no external torque in the ATP synthesis mode (**A-C**), a weak opposing torque 8.6 pN · nm in the ATP synthesis mode (**D-F**), and a strong torque 86.2 pN · nm in the ATP hydrolysis mode (**G-I**). Trajectories for the WT are shown in gray for comparison. (**C, F, I**) The average number of transferred protons per revolution are plotted for each type of transports: From the left to the right, transfer from IMS to matrix side, pump from matrix to IMS side, leak from IMS to matrix side, "leak" from matrix to IMS, slip from matrix to matrix, slip from IMS to IMS.

Interestingly, in the aR176A, the most frequent proton transportation pathway was Path 2, rather than Path 1, the dominant pathway in WT (Fig.S3). Namely, from DEP(c_I_, c_J_), the protonation at c_I_ occurs before deprotonation at c_A_. The resulting DEP(c_J_) state, which corresponds to an intermediate state of Path 2, becomes a broader distribution in rotary angles, which results in increase in the population of this intermediate state and the Path 2. These changes can be understood in the following manner. Compared to the WT case, because the attractive interaction between deprotonated cE59 and aR176 disappears in the aR176A case, the energy barrier on protonation of c_I_, the bottleneck in Path2, disappears. This results in the acceleration of Path 2. We also note that, compared to the WT, the angle distribution in the initial DEP(c_I_, c_J_) state of the aR176A moved downward, i.e., clockwise rotated (Fig.S3), which increases the energy barrier for the first step of Path 1. On top, the lack of the attractive interaction slows down the rotation of F_O_ due to the lack of driving force for the counterclockwise rotation.

Broadening of the angle distribution in DEP(c_J_) of the aR176A mutant increases the proton leakage as well. We denote the proton leakage path observed in aR176A without external torque as Path 7,

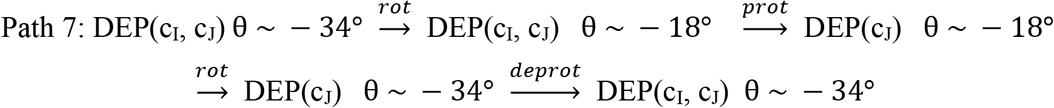

In the path, the lack of electrostatic attraction of aR176 also contributes to the clockwise rotation 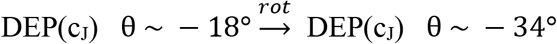.

Next, we tested a sensitivity of the mutant with respect to a weak opposing torque, which could mimic the load by F_1_. With a constant and weak torque 8.6 pN · nm, we repeated the otherwise same MC/MD simulations. The results (Fig.4D) show qualitatively similar behavior with a more enhanced difference. While the WT F_O_ rotated nearly at the same angle velocity as the case of no torque, the aR176A mutant exhibited more stochastic and, on average, slower rotation (Fig.4DF).

### The aR176A does not have the proton pump function

Next, we applied a strong external torque, which corresponds to the ATP hydrolysis mode, to the aR176A mutant F_O_. While the torque-driven rotation velocity is nearly the same between WT and the aR176A, there is a critical difference in the proton pump function between the WT and the aR176A (Fig.4G). The proton leakage drastically increased in the aR176A compared to the WT, and the aR176A pumps almost no proton in total (Fig.4GI). It has already been assumed that the mutation at the corresponding arginine in a bacterial F_O_F_1_ lacks proton pump activity (*29*) and our result supports this assumption. From our result, it turned out that actually aR176A transports some protons from the matrix to the IMS sides, but it leaks almost the same number of protons towards the opposite direction, resulting in almost no net proton pumping.

Since the lack of aR176, which means that no barrier between IMS and c-subunit that locates near matrix side, protons can enter into c_J_ from IMS side directly before c_A_ receives a proton from matrix. Because c_J_ locates near at the matrix side, a proton they have can leave from c10-ring and exit to the matrix side.

### The c_J_E59A mutant rotates with reduce velocity and power

We also performed hybrid MC/MD simulations for the mutant in which one of the cE59 is substituted with alanine (c_J_E59A). Without the external torque, the c_J_E59A mutant can rotate with a reduced velocity (Fig.4B). The plot shows that the rotation pauses every 360°, apparently corresponding to the mutant stoichiometry of the c_10_-ring. Additionally, the number of protons transported per revolution is about 9 (Fig.4C), which is a reasonable number because the c_J_E59A lacks one protonatable site out of 10.

Although it is experimentally confirmed that the cE59A mutant loses its ATP synthesis activity, it successfully transports protons from IMS to matrix and rotates in counterclockwise direction in our model. It is because our model lacks F_1_. In reality, F_O_ protein is connected to the central rotor to conduct its torque to F_1_, hence it receives some resisting power to induce conformational changes in F_1_. In order to take this into account, we applied a small torque, 8.6 pN · nm and repeated the otherwise same simulations (Fig.4E). Fig.4E shows that the c_J_E59A mutant hardly rotated, although WT rotated nearly equally to the case of no load (gray in Fig.4BE). This result suggests that the c_J_E59A mutant is less powerful than the WT, i.e. the mutation makes it impossible to generate a torque strong enough to drive F_1_ to synthesize ATP.

### The c_J_E59A mutant works as a proton pump, though it is less efficient

At last, we applied a strong torque 86.2 pN · nm to the c_J_E59A that corresponds to the ATP hydrolysis mode. The cumulative rotation angles are plotted in Fig.4H. As shown also in Fig.4HI, it works as a proton pump though it is less efficient. Since it loses one component of c_10_-ring, the number of protons transported decreases compared to the WT.

### The dependency of the rotary mechanism on the computational parameters, pKa

So far, we used one set of pKa values in 12 glutamates based on previous suggestions (*20*). However, these values are either merely assumed or inferred from indirect data; there is no direct measurement on these pKa values. Here, to investigate how F_O_ functions can differ depending on pKa values in key glutamates. Here, we changed pKa values of aE162 as in Table S2 and repeated otherwise the same simulations; we performed simulations with no load (as the ATP synthesis mode) and with a strong load, 86.2 pN · nm (as the ATP hydrolysis mode).

Results in Fig.5 show that the pKa value of the aE162 does not strongly affect the rotational velocity and the rate of proton transfer in the ATP synthesis mode (Fig.5ABC). With a smaller pKa value of aE162, the protonated aE162 is less stable, which slows down the proton transfer from cE59 to aE162, but accelerates the subsequent release to matrix aqueous environment by the same factor; the two effects cancel each other. The similar argument could be applied to the pKa value of aE223.

**Fig. 5.**
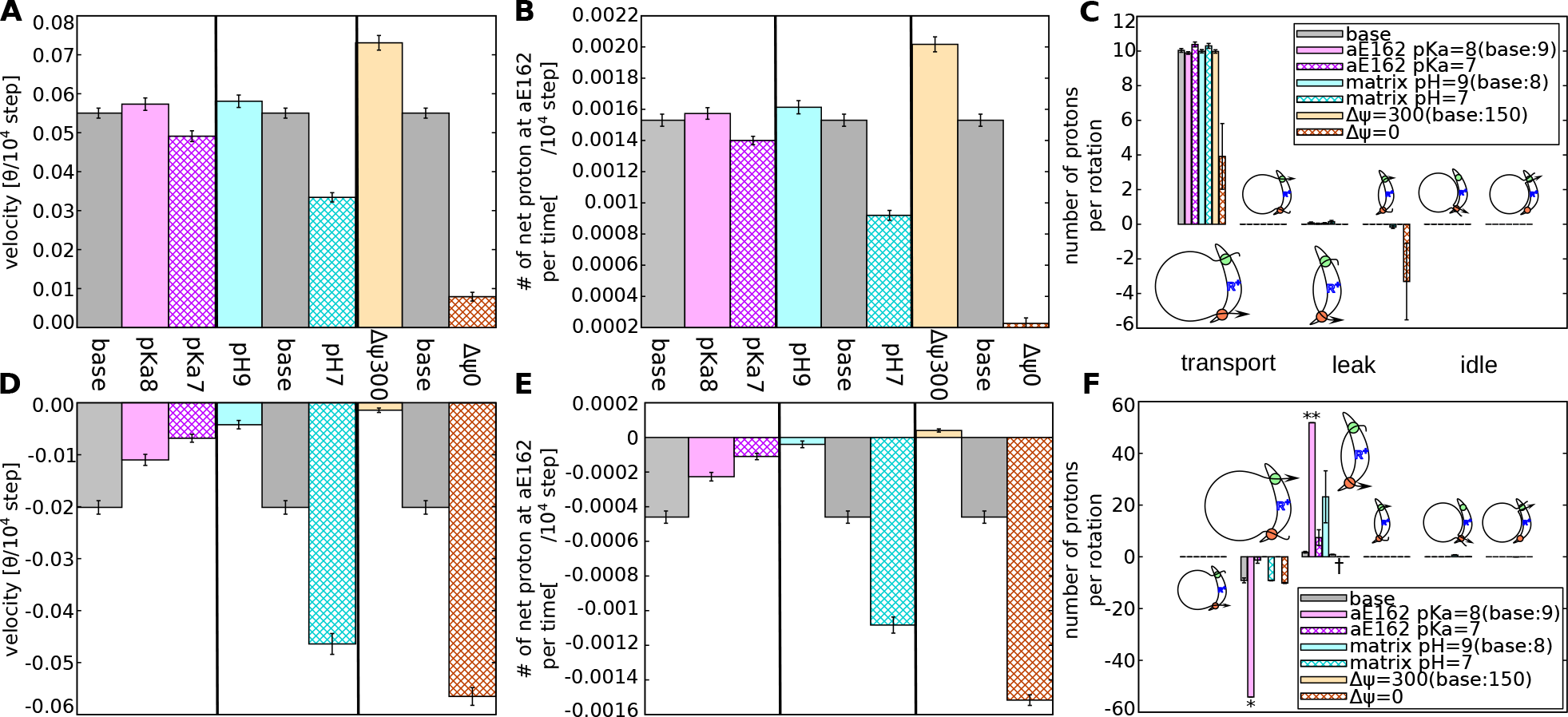
The effect of the parameters, pKa, pH, ΔΨ on the model F_O_ functions. The rotational velocities, numbers of net proton movement per time and per rotation, are plotted with no external torque in the ATP synthesis mode (**A-C**) and a strong torque 86.2 pN · nm in the ATP hydrolysis mode (**D-F**). The pKa value does not strongly affect both the rotational velocities and the number of transferred protons in the ATP synthesis mode (**A-C**) but decreases that of the ATP hydrolysis mode (**D-F**). The pH and ΔΨ differences affect in the expected way. With strong proton motive force, ATP synthesis activity increases and proton pumping activity is inhibited. Note that the value indicated with “*”, “**”, and “†” means 42.3, 42.5, and N/A, respectively.

On the other hand, in the ATP hydrolysis mode, the pKa value of aE162 has sharp effects. A smaller pKa value, 8 or 7, decreases the rotational velocity and sharply reduces the rate of proton pump (Fig.5DE). In the ATP hydrolysis, a proton must be taken in from the matrix side via aE162. A smaller pKa value of aE162 makes the proton uptake from the matrix side more difficult, thus the rotational velocity decreases. This process is a kinetic bottleneck in the ATP hydrolysis mode.

In the ATP hydrolysis mode, a proton entered into c_10_-ring from IMS side, instead of the matrix side, causes proton leakage (see Path 6). Generally, proton concentration in the matrix side is lower than the IMS side so that proton uptake from the matrix side is slower than that from the IMS side. Therefore, it is hard to strictly stop the proton leakage while keeping the ATP synthesis ability. Actually, as the pKa of aE162 decreases, the probability of aE162 having a proton also decreases and it increases the proton leakage from IMS to matrix (Fig.5F). With the value 7.0, the number of protons leaked from IMS to matrix is about the same as the number of protons pumped from matrix to IMS. Thus, with that value, F_O_ can virtually pump up proton. Those results show that the range of parameters that allows F_O_ to work as both ATP synthase and proton pump is narrow and our parameter set used here is inside of it.

### The effect of pH

Next, we address how pH in the IMS and matrix environments affect the F_O_ function; notably, the difference in pH’s, but not individual pH’s, modulates the dynamics. Here, we varied the pH value in the matrix side (see Table S2) and performed otherwise the same simulations as above. Results in Fig.5 show that a larger value of the pH in the matrix side makes the rotational velocity and the rate of proton transfer larger in the ATP synthesis mode (Fig.5ABC), as expected from experimental results (*30*). The large pH of matrix side facilitates proton release from aE162 and thus the proton transfer from c_10_-ring to aE162 is more efficient. As discussed before, the proton transfer at the matrix side often becomes a rate-limiting process; hence it is reasonable that a large pH in the matrix side increases the rotational velocity in the ATP synthesis mode.

On the contrary, in the ATP hydrolysis mode, a smaller pH value in the matrix side makes the rotational velocity and the rate of proton pumps larger (Fig.5DEF). The reason coincides with the case of ATP synthesis process. When the matrix side pH becomes larger, the probability of aE162 being protonated becomes smaller, thus the proton transfer from aE162 to c_10_-ring is enhanced. At the same time, proton leakage also occurred via the same mechanism as the case of a large pKa value at aE162 because the probability of being protonated is controlled by the difference between pKa and pH.

### The effect of ΔΨ

Finally, we investigate the effect of the membrane potential; Either deleting or doubling the membrane potential (Table S2), we repeated the simulations as before. In the ATP synthesis mode, a larger membrane potential makes the rotational velocity larger in the ATP synthesis process (Fig.5ABC), as expected from experimental results (*30*). It is because the protons are stabilized by being passed to aE223 from IMS and to matrix from aE126.

On the other hand, in the ATP hydrolysis mode, a smaller membrane potential makes the rate of proton pumps larger (Fig.5DEF). This is because the energy cost to make aE162 protonated decreases. At the same time, the possibility of aE223 being protonated decreases and thus the proton leakage become less probable. It drastically accelerates the proton pumping.

Notably, within our simulation method, both pH and ΔΨ values appear only in the same formula that represent the probability of protonation at the glutamates in the two half channels (see Materials and Methods: Simulating Proton-Transfer Coupled Protein Dynamics). Thus, the total proton motive force, but not pH and ΔΨ separately, controls the entire behaviors, which is experimentally well examined (*30*).

### Protonation state dependent free energy surfaces

In order to investigate why the Path1 is preferred than the Path 2 and the Path 3, we estimated free energy surfaces along the rotation angle for different protonation states (see more detailed method in Section S1). In this analysis, to compare the free energies, we need to keep the total number of protons in the system the same. Thus, for the Path 1/3 and Path2, respectively, we include 9 and 10 protons in the system and explored the free energy surfaces separately (Fig.6AB).

**Fig. 6.**
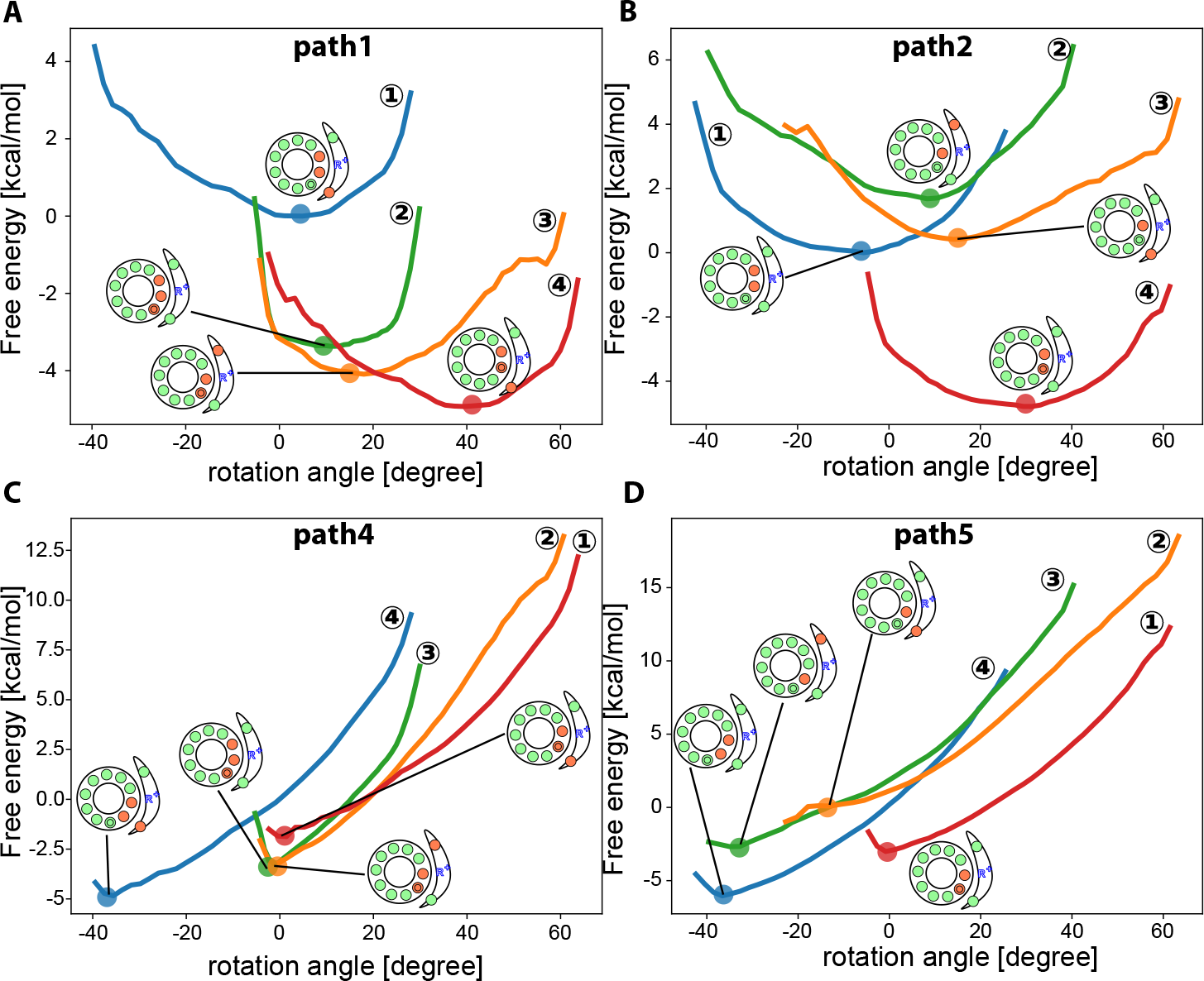
Free energy curves of individual protonation states. **(A)** Paths 1 and 3 transit four free energy surfaces in the following order; DEP(c_I_, c_J_; aE162) in blue, DEP(c_I_, c_J_, c_A_) in yellow, DEP(c_J_, c_A_; aE223) in green, and DEP(c_J_, c_A_; aE162). **(B)** Path 2 goes through four free energy surfaces in the order of DEP(c_I_, c_J_) in blue, DEP(c_J_; aE223) in yellow, DEP(c_J_; aE162) in green, and DEP(c_J_, c_A_) in red. **(C-D)** In C and D, the external torque of 86.2 pN · nm is applied to the data in A and B, respectively.

Figure 6A shows free energy surfaces without the external torque. The first transition in the protonation states in Path1 drastically decreases the free energy compared to that of Path2 (Fig.6B). Additionally, after the protonation of c_I_, the most stable angle is almost kept, suggesting that the significant rotation occurs after exchanging protons with bulk. The significant rotation might possible before the protonation at c_I_, but it has large energy barrier. This is why Path 1 is more preferred than Path 3. On the other hand, the first protonation state transition in Path 2 slightly increases the energy. This is the reason why Path1 is the dominant pathway compared to Path 2.

### Estimation of the torque generated by the F_O_ motor

In order to synthesize ATP using proton motive force, F_O_ needs to generate a torque strong enough to induce conformational change in F_1_. Therefore, how much torque c_10_-ring generates by the proton motive force is important.

To estimate the torque generated by the F_O_, as a function of the external torque to the F_O_ we tested if our model F_O_ can rotate in the direction of ATP synthesis or not by the default proton motive force. The 86.2 pN · nm of torque resulted in the reverse rotation (ATP hydrolysis direction), coupled with the proton pumping, as described. Thus, here we test weaker torques varying between 0 and 86.2 pN · nm. F_O_ rotated in the ATP synthesis direction up to 43.0 pN · nm of torque transporting protons from IMS to matrix. Whereas, with the 51.6 pN · nm of torque, we observed, as the average, the reverse rotation (Fig.S4). An external torque at which the net rotation becomes zero, which corresponds to the torque generated by F_O_, is estimated as ~50 pN · nm. Interestingly, our result coincides with the previous theoretical prediction by Oster (*31*), from 40 to 50 pN · nm.

From this result, we can estimate the efficiency of the energy conversion from proton gradient to the torque by our F_O_ model. Since c_10_-ring rotates in 36° per one proton, the energy converted to torque per one proton is

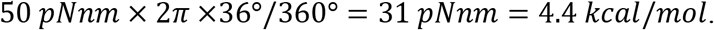

On the other hand, the free energy loss due to one proton transfer from the IMS to the matrix side is a sum of the energy owing to the difference in pH and the membrane potential. The former is

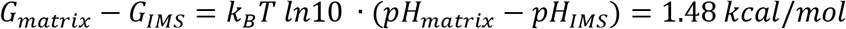

when the pH in IMS and matrix are 7 and 8, respectively. When the membrane potential is 150 mV, the corresponding energy is simply 0.15/0.04337 = 3.46 *kcal*/*mol*. Thus, the total free energy consumption per one proton is 1.48 + 3.46 ~ 4.94 *kcal*/*mol*.

Taken together, the maximum efficiency in free energy transduction of our model F_O_ motor *η* is estimated as *η*~89 %.

### The effect of membrane region

As discussed so far, the location of the protein-membrane interface near the matrix channel has an important role for the rotary movement in both ATP-synthesis and proton pumping activity. It is known that mutations in aromatic residues around the matrix channel causes mitochondrial diseases (*32*), possibly because the mutation makes a-subunit be distant from c-ring and the lipids intrudes into the gap. Here we investigated this effect by changing the position of the surface of the implicit membrane potential *V*_*mem*_ (Fig.S5A). We performed the hybrid MC/MD simulations in the ATP synthesis mode, with varying angles of borders between lipid phase and a-subunit-facing phase; the border is changed from the angle of WT to the angle at aE162 (Fig.S5A).

Interestingly, the rotational velocity of the c-ring decreases monotonically (Fig.S5B) and the number of protons transferred per rotation also decreases (Fig.S5C). This is caused by the wider membrane region. Since the protonated state is more preferable in the membrane, in order to pass a proton to the matrix channel, cE59 needs to escape from the membrane region solely by the diffusion. This is similar to the case of c_J_E59A and the number of transported protons is almost the same as that of c_J_E59A (Fig.5C).

Especially, when the membrane region locates at the aE162, no protons are transported but protons leak without rotating the c-ring (Fig.S5C). In this case, passing a proton increases the energy when the cE59 approaching from the membrane region but it does not when the cE59 approaches in the opposite direction. As the proton transfer becomes difficult, the leaky pathway becomes more popular. The mechanism is similar to the case of the aR176A mutant, in which protons leak without rotating c-ring.

The result explains why the mutations in the aromatic residues around the matrix side result in loss of its activity. As the a-subunit become distant from c-ring, the membrane-protein interface invades toward the matrix channel and it affects the activity in similar ways to the cases of aR176A and c_J_E59A.

## Discussions and Conclusion

The current model F_O_ motor reproduced the rotations in the ATP synthesis direction coupled with proton transfers across the membrane driven by the proton motive force as well as the proton pump functions coupled with the reverse rotations when the torque was applied in the ATP hydrolysis direction. Furthermore, the inactivation of ATP synthesis ability and/or proton pump ability for mutants was also reproduced. By visualizing the movement of every protons in time domain, we found that protons-transfer from c_10_-ring to a-subunit before proton-transfer from a-subunit to c_10_-ring is the dominant pathway in both synthesis and hydrolysis modes.

In the ATP synthesis mode, the de-protonation of c_10_-ring cE59 plays a central role in rotational movement, while in the ATP hydrolysis, the rate-limiting event is the protonation of c_10_-ring cE59. It is clear that proton transfers occur efficiently when the donor and the acceptor are close enough, but, in addition, the range at which c-ring faces the lipid membrane is a crucial factor to determine the pathways. Within the lipid-facing membrane region, cE59 is hardly deprotonated, due to the electrostatics in the hydrophobic lipid regions. Therefore, it may be interesting to investigate the detailed pathways of proton-transfer coupled rotations for F_O_ motors from different species with different numbers of c-subunits.

### Limitations of this work

Our current F_O_ model is a minimal model to address proton-transfer coupled F_O_ rotations at structural details. First, for computational ease, we utilized a coarse-grained model throughout, which is lower resolution and unavoidably less accurate than the gold-standard atomistic molecular mechanics. The similar proton-transfer coupled atomistic MD is highly desired in future. Second, our model contains only a-subunit and c_10_-ring. However, in the actual F_O_ motor, additional subunits such as b-subunit exist. If these subunits play important roles other than just stabilization of the relative positioning, the ignorance of them could affect the motions/pathways. In addition, because F_1_ has 3-fold symmetry, being mismatched to the 10-fold symmetry in c_10_-ring, the whole F_O_F_1_ ATP synthase must show more complex and asymmetric behaviors, which can be a future direction to investigate.

## Materials and Methods

### Experimental Design

Fig.1 shows the overview of the simulation setup used in our research. Fig.1BC shows the F_O_ ac_10_ complex used in our simulation. Fig.1D schematically describes the hybrid MC/MD simulation that alternatively update the protein configuration by MD and the protonation state by MC. More precise methods are described in the following three sections.

### Protonation-state dependent Hamiltonian

We first describe a general formulation that concisely represents the protonation-state dependent protein energy. We assume that the target protein contains *n*_*p*_ amino acids that can take both protonated and deprotonated states, which we call the protonatable sites. We introduce an *n*_*p*_-dimensional vector 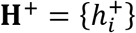, of which each element 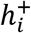 represents the state of the protonatable site and takes either 1 when protonated, or 0 when deprotonated. Then, we introduce a protonation-state dependent Hamiltonian for proteins in general,

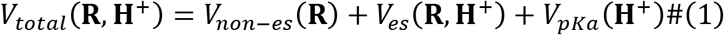

where **R** collectively represents coordinates of the protein structure. The first term in the right-hand side is the protein interaction energy excluding the electrostatic interactions. The second term represents the electrostatic interaction that depends both on the structure **R** and on the protonation state **H**^+^. The last term is for the intrinsic energy for the protonation of individual protonatable amino acids, which depend on their intrinsic pKa values at given environment.

### Protein Modeling

Our simulation system contains an a-subunit and 10 c-subunits that form the c_10_-ring. The structure model is based on the cryo-EM structure of a yeast mitochondrial ATP synthase (PDB ID: 6CP6(*20*)) (Fig.1A). For the c_10_-ring structure, we first trimmed one C-terminal residue in two chains to make the chain length uniform (chains M and R in the PDB file). While the c_10_-ring structure in the cryo-EM model is asymmetric due to the interaction with other subunits, especially with a-subunit, we need a ground state conformation of the c-subunit alone. As a proxy, we obtained the c-subunit structure averaged over all the 10 c-subunits in 6CP6, which was used as the reference structure of the c_10_-ring. For all the proteins, each amino acid is coarse-grained to a single bead located at the position of the Cα atom of the residue. Membrane and water solvent are treated implicitly via effective potentials.

For *V*_*non*−*es*_ (**R**) in eq.(1), we used the AICG2+ potential for the intra-chain interactions of a-subunit, c-subunit, and for the interface interactions between neighboring c-subunits (*25*, *26*), where the energy function is biased to the experimental reference structure and several energy parameters are tuned based on all-atom force field energy at the reference structure. Between a-subunit and c_10_-ring, we included only the excluded volume interactions in *V*_*non*−*es*_ (**R**), but not the structure-based interactions, in order not to inhibit the rotation.

The electrostatic term in eq. (1), *V*_*es*_(**R**, **H**^+^), consists of two contributions; the standard Coulomb interaction between any charged residues V_*c*_(**R**, **H**^+^) and the membrane-environment term V_*mem*_(**R**, **H**^+^). The proteins have both fixed charges and charges on the protonatable sites. The latter contains 12 glutamic acids according to Srivastava et al.(*20*)(Fig.1BC); cE59 in each of 10 c-subunits, aE223 and aE162 in the a-subunit. The aE223 resides at the interface to c-ring and connects to the IMS side via the IMS half channel, whereas aE162 sits at the interface to c-ring connecting to the matrix side. We assign −1 charge for the deprotonated state and 0 for the protonated state of these 12 glutamic acids. For other fixed charged residues, we put +1 charge on arginine and lysine residues and −1 charge on aspartic acid and glutamic acid residues. We set the dielectric constant of the system to 10 considering hydrophobic membrane environment. For mutants, we changed the residue type and the charge on it.

The membrane-environment term approximates a free energy cost when a charged amino acid is embedded into the central hydrophobic layer of lipid membrane. Specifically, cE59 sits in the middle of the transmembrane helix, facing outward of the ring form. When the deprotonated cE59 faces directly to lipid, we assign a constant free energy cost, *ϵ*_*mem*_. On the other hand, when the deprotonated cE59 is at the interface to a-subunit, no free energy cost is applied. Thus, the free energy value for deprotonated cE59 differs between at the interface to lipid and at the interface to a-subunit. We model this difference in the free energy as a smooth and implicit membrane-environment potential applied to charged residues.

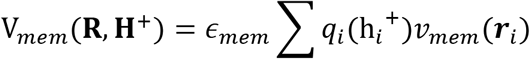

where *q*_*i*_(h_*i*_^+^) = h_*i*_^+^ − 1 represents the charge of cE59 and the potential 𝑣_*mem*_(***r***_*i*_) is a smooth function that takes 1 when the residue *i* faces directly to membrane lipid and switches off at the interface to a-subunit (Fig.S1A). We note that the protonated, and thus uncharged, cE59 has no such energy cost. Specifically, to define 𝑣_*mem*_(***r***_*i*_), we first define the two border planes (dashed lines in Fig. 1C with the angle 51.2 and −69.6 degree from x-axis) that separate the lipid-facing region (gray shaded in Fig.1C) and the a-subunit facing region (white in Fig.1C). For each of the borders, we then define a coordinate *x*(***r***) normal to the border plane and that takes the positive value in the lipid membrane side. We employed a smoothly switching function,

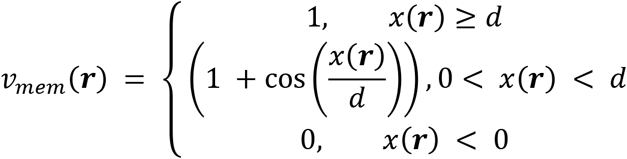

The free energy cost *ϵ*_*mem*_ was set as 10 [kcal/mol]. The *d* is a control parameter to modulate the breadth of the border and is set as 0.4 [nm].

In the full F_O_F_1_ system, the c_10_-ring would be tightly attached to the central rotor of the F_1_ subunit to transmit the torque generated by the rotation of c_10_-ring. These interactions together with the peripheral stalk should keep the c_10_-ring close to the a-subunit and, at the same time, also allow it to rotate. Whereas, in our simulation setup, we only have a- and c-subunits. Thus, in order to prevent the c_10_-ring from tilting and diffusing away from a-subunit, we added an extra-restraint that keeps the rotational axis of the ring to the z-axis (Fig.1C). To define the rotational axis, we choose two residues from each c-subunit that locates the first and third quarter in z coordinate and fit a circle to them. Then we applied harmonic restraints to the residues to keep the distance between them and the center of the corresponding circle. We also applied 3 harmonic restraints to the a-subunit to prevent free diffusion and rotation after aligning aR176 on the x-axis.

The last term in eq. (1), *V*_*pKa*_(**H**^+^), simply reflects the intrinsic preference of the protonation at each site and can be written as,

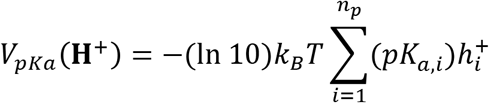

where *pK*_*a,i*_ takes into account the intrinsic pK_0_ of glutamic acid as well as its basal environment in Fo motor. Specifically, we set pK_*a*_(*cE*59) = 8.0, pK_*a*_(*aE*223) = 6.0, and pK*a*(*aE*162) = 9.0, as the default values (*20*), while we will vary these values later to test roles of these values. We note that, as shown in Fig.1A as red and green beads, all the 12 protonatable sites locate almost the same z-coordinate values in the membrane. Thus, we set the membrane potential of all the protonatable sites identical and thus the membrane potential does not appear in the Hamiltonian. Instead, the membrane potential appears in the boundary condition, as described below.

To identify each subunit in the c_10_-ring, we assign an alphabet “A” to “J” to each subunit in the clockwise manner looking from F_1_-side (Fig.1B). Note that these alphabets are associated with molecules; when the c_10_-ring rotates, these alphabets rotate together.

### Simulating Proton-Transfer Coupled Protein Dynamics

Given the protonation-state dependent Hamiltonian, we now describe the dynamic simulation method. In the current modeling, a proton transfer can be modeled as a change of a pair of charges on two protonatable sites; the proton donor changes its charge from 0 to −1, while the proton acceptor changes its charge from −1 to 0. Because traditional MD simulation is not capable to treat these charge changes, we combined the MD simulation with the MC transitions, which represent proton transfers, to realize the proton-transfer coupled rotation of F_O_ *in silico*. Short-time MD stages and MC stages are conducted alternatively as in (*33*), which is also of some similarity to constant pH MD (*34*, *35*). In the MD stage the protein structure moves with a fixed protonation state, whereas in the MC stage the protonation state changes with a fixed protein structure (Fig.1D).

In each MD stage, with the given protonation state, we perform a MD run for *τ* = 10^5^ MD steps (Fig.1D). The MD simulations were conducted by the underdamped Langevin dynamics at a temperature of 323K. We used default parameters in CafeMol except the friction coefficient, which is set as 2.0 to model the effect of membrane environment. All MD simulations are performed using the CafeMol package version 2.1(*36*).

In the MC stage, we first determine the protonation states of the IMS-connected site aE223 and the matrix-connected site aE162. We assume that the aE223 and aE162 sites are able to exchange protons with the IMS and matrix aqueous environments, respectively, and that the proton exchange between these sites (aE223 and aE162) and the aqueous environment is sufficiently fast compared to the timescales of F_O_ rotational motions. Hence, the protonation states on these sites are well equilibrated with the respective environment based on the pKa, the environmental pH, and the membrane potential ΔΨ. Unless otherwise denoted, we set the pH values of IMS and matrix as 7.0 and 8.0, respectively, and the mitochondria membrane potential as ΔΨ = 150mV; the membrane potential at the matrix environment is −150 mV relative to the IMS environment. Since the 12 protonatable sites are located near the center of the membrane, we set the membrane potential of all these sites as ΔΨ/2. Given that, we obtain the local equilibrium probabilities of protonation at aE223 and aE162 as,

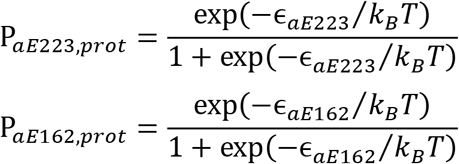

where

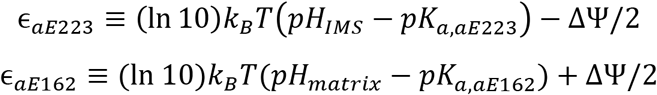

There protonation probabilities serve as the boundary conditions. Each MC stage begins with sampling the protonation state of these two sites, aE223 and aE162, from these equilibrium probabilities.

Once the protonation states of aE223 and aE162 are set in each MC stage, we then simulate proton transfers among the 12 protonatable sites. Each proton in the protonated site can transfer to a deprotonated site, keeping the total number of protons unchanged. We limit the transfer to be between c-subunit sites and a-subunit sites. For the *i*-th proton, its transfer probability to the *j*-th site depends on the kinetic weight factor *w*_*j*←*i*_ and the energy change upon the proton transfer Δ*E*,

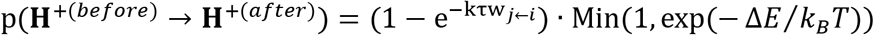

where **H**^+(*after*)^ and **H**^+(*before*)^ represent the protonation state after and before the trial of a proton transfer, respectively, and *k* is the overall rate constant and τ = 10^5^ MD steps.

The kinetic weight factor *w*_*j*←*i*_ reflects the geometry of the donor and acceptor residues. The weight is calculated by a function that depends on the distance *r* between the donor and acceptor sites as well as an angle *θ* between two vectors that emulates the conformation of side chain (see Fig.S1B). This modeling is similar to the recent coarse-grained modeling in DNA and RNA base-base interactions(*37*, *38*). The first vector connects the protonation site on c_10_-ring with the site on a-subunit. The second vector connects the middle point of the adjacent residues on the c_10_-ring and the site on c_10_-ring, which indirectly monitors the side chain orientations (Fig.S1B). The functional form is given as,

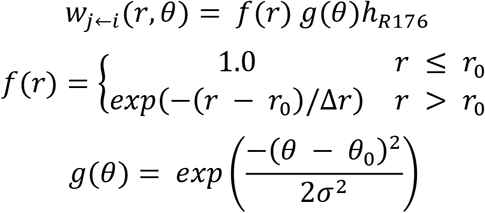

The *f* is an exponentially decaying function of the distance with the threshold distance r _0_ = 0.8nm at which the tip of the side chain of glutamic acids are nearly in the direct contact and Δ*r* = 0.4 *nm*. The *g* is a Gaussian function that has the peak at *θ*_*0*_, the corresponding angle in the reference cryo-EM structure, and the width σ = 36°. Notably, the above defined geometric factor *w*_*j*←*i*_ is symmetric with respect to **H**^+(*before*)^ and **H**^+(*after*)^ so that the MC algorithm, together with the Metropolis criterion, satisfies the detailed balance. Given that we do not know the absolute overall rate *k*, we simply chose it as *kτ* = 1.

The trial proton transfer is accepted by the Metropolis criterion,

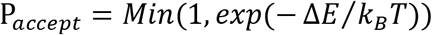

with

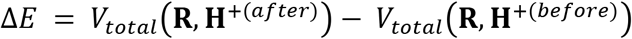

or rejected otherwise. Notably, protein structures do not change in this stage and thus *V*_*non*−*es*_(**R**) would not contribute in the energy change.

Additionally, we introduced an extra geometric factor *h*_*R*176_. It is known that the arginine in a-subunit, aR176, serve to block proton leaks from the IMS to the matrix environment without transferring into c-subunits, which thus separates two half-channels of the a-subunit (*21*, *39*). In order to take this effect into account, we put a simple factor *h*_*R*176_, which is normally unity, but takes zero when the y-coordinates of the aR176 is in between the y-coordinates of the proton donor and acceptor sites (see Fig.1AB for the definition of the coordinates). For the simulation of the mutant on aR176, we removed this extra factor.

In each MC stage, we repeat these MC trials for every proton in protonatable sites in random order. At the first stage of the simulation, we conducted one round of a MD stage for the equilibration and then we performed 3000 rounds of the hybrid MC/MD simulations.

## H2: Supplementary Materials

Section S1. Analyzing Free Energy Differences Between Different Protonation States

Fig. S1. Simulation setup

Fig. S2. The trajectories of the WT with non-zero proton-motive-force in the ATP hydrolysis mode

Fig. S3. Comparison of the mechanisms between aR176A mutant and the WT in the ATP synthesis mode

Fig. S4. The number of revolutions per trajectory under different external torque.

Fig. S5. The effect of the lipid membrane border on the F_O_ rotation

Table S1. The number of proton-transfer per rotation

Table S2. The list for Fig.5

Movie S1. ATP synthase mode

Movie S2. ATP hydrolysis mode

## Author contributions

S.K. and S.T. conceived the study. S.K., T.N., and S. T. designed the study. S.K. and T.N. developed the simulation code, performed simulations, and analyzed the data. All the authors discussed the results and wrote the paper.

## Data and materials availability

## Supplementary Materials

### Section S1. Analyzing Free Energy Differences Between Different Protonation States

We compared the free energy curves in different protonation states. We chose two representative pathways of proton-transfer coupled rotary motions observed in our MC/MD simulation described above and picked 8 protonation states from the pathways. Then we carried out 10 independent 10^8^ MD simulations for each state with 8 fixed protonation states. Since we need to fix the states of each protonatable site while the MD simulation, the states of aE223 and aE162 are fixed in this simulation. We chose the protonation states of the a-subunit to make the total number of protons kept upon the transitions along each pathway.

Using those samples, we employed the Bennett acceptance ratio method (*40*) and calculated free energy differences between the states that are next to each other in the pathways. For this calculation, we applied the protonation-state dependent Hamiltonian. Applying the method to each pair of states, we evaluated the free energy differences of each transition in the pathways. Because the effect of proton exchanges between the a-subunit and bulk are not considered in the procedure, the net free energy differences after the step of 36 degrees of rotation became zero in each proton transfer pathways, confirming the accuracy of our calculations. To consider the stabilization of transmembrane proton transfer, we reinterpreted the proton movement in the step in which a proton is exchanged between aE223 and aE162 as proton exchange between each residue and the bulk. As a result, we obtained the free energy differences between the first state and the state after 36 degrees of rotation as −4.9 kcal/mol that is the same value as the stabilization by one proton transfer from IMS to matrix in our standard setup.

**Fig. S1.**
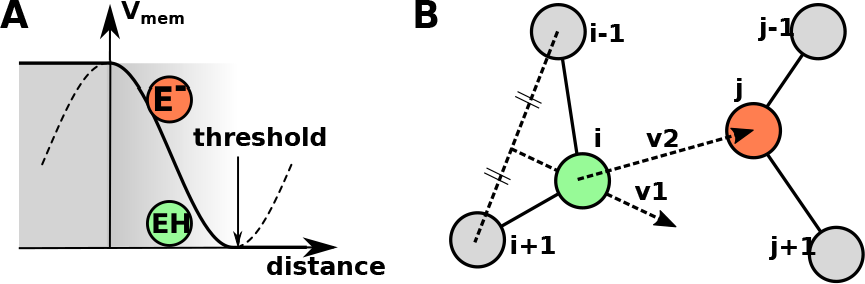
Details in the simulation setup. **(A)** The schematic view of our implicit membrane potential, V_*mem*_(**R,H**^+^). We applied a free energy cost to the charged, i.e., deprotonated residues. If the residues are protonated, the free energy cost vanishes. **(B)** The illustration of the geometric parameters used in the kinetic weigh *w*. The first vector 𝑣_1_ connects the middle point of the adjacent residues on the c_10_-ring (left gray circles) and the protonation site on c_10_-ring (green). The second vector 𝑣_2_ connects the protonation site on c_10_-ring (green) and the site on a-subunit (red).

**Fig. S2.**
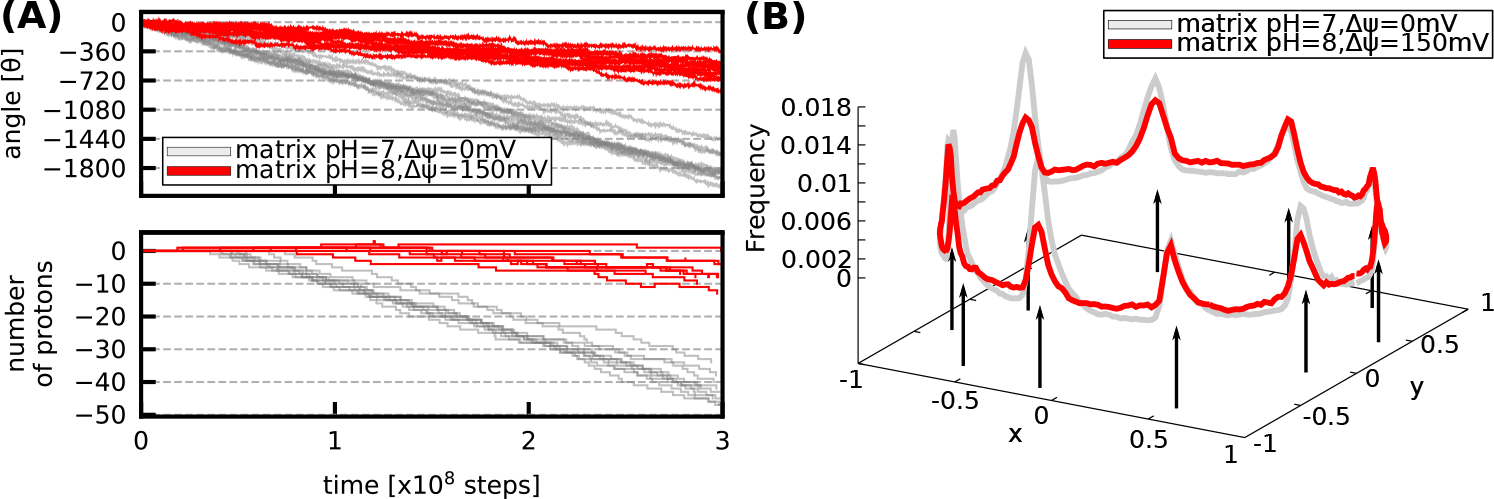
The trajectories of the WT F_O_ with non-zero proton-motive-force in the ATP hydrolysis mode. **(A)** Time courses of the cumulative rotation angle (top) and the number of protons that across all the way from IMS side to matrix side (bottom) for 10 trajectories. The negative rotation angle means the clockwise rotation corresponding to proton pumping. Note that the negative number of protons means the protons transfer in the opposite way, from matrix side to IMS side. **(B)** Probability distribution of the rotation angle in the representative trajectory.

**Fig. S3.**
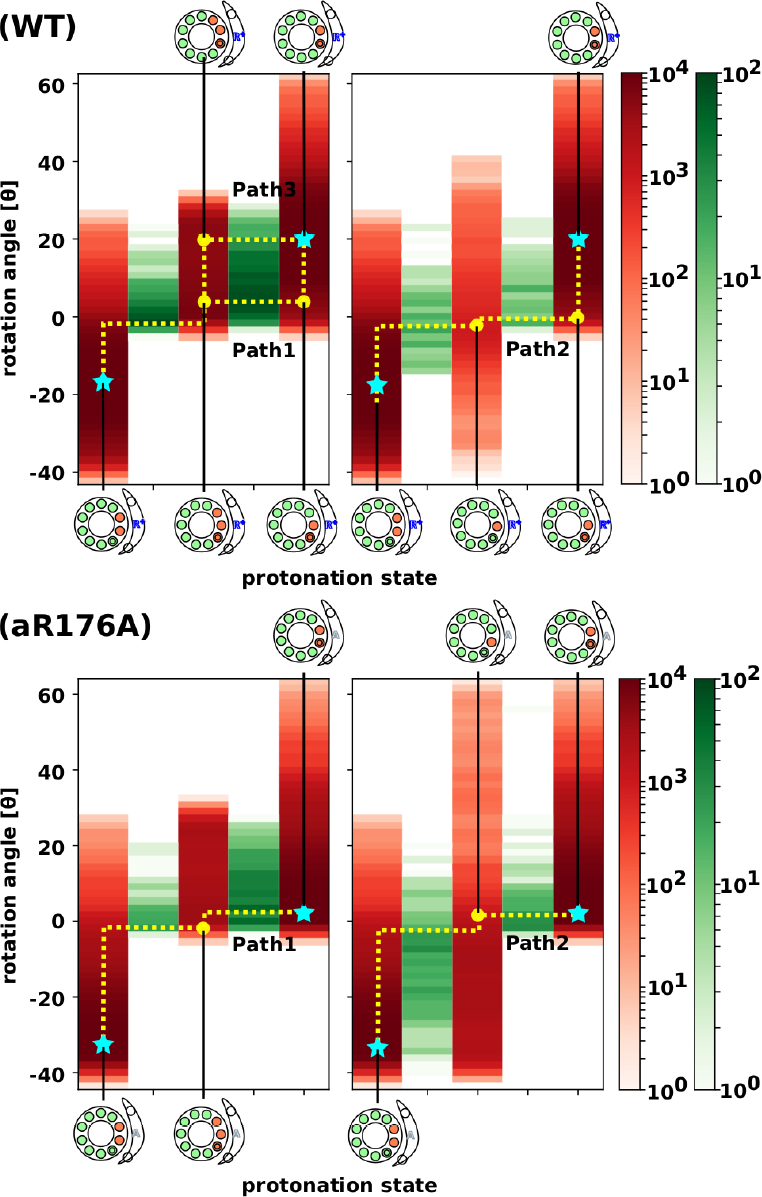
Comparison of the mechanisms between the aR176A mutant and the WT F_O_’s in the ATP synthesis mode. **(WT)** The same as the one shown in Fig.2D. **(aR176A)** The result of the same analysis for the aR176A mutant.

**Fig. S4.**
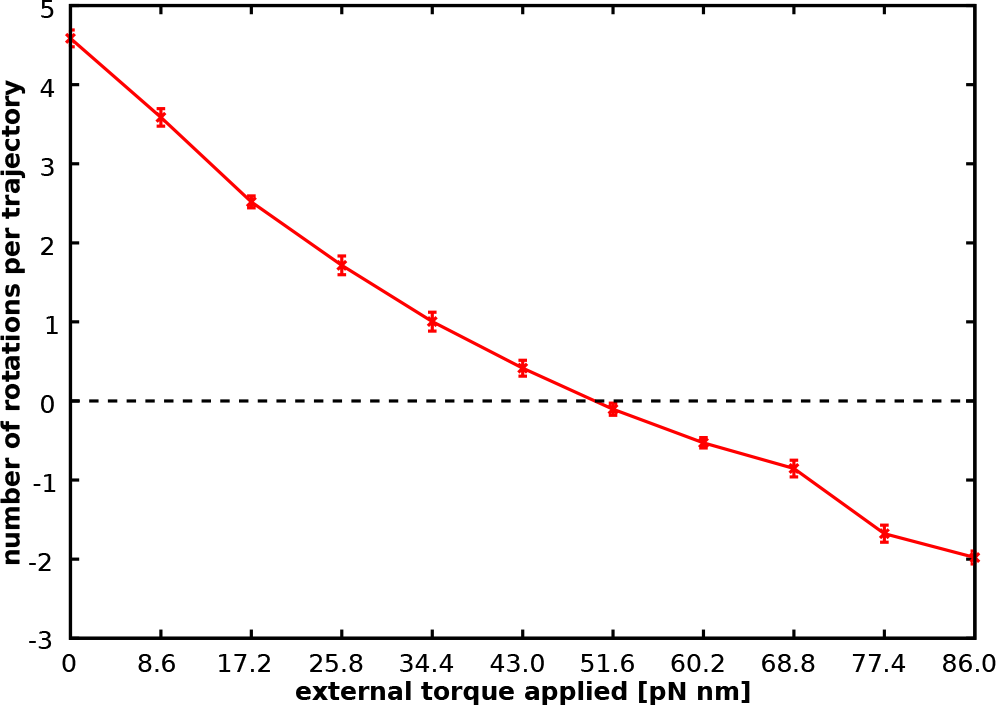
The number of revolutions per trajectory under different external torque. The number of revolutions in one trajectory is plotted. Between 43.0 pN · nm and 51.6 pN · nm, the number of rotations turned from positive to negative.

**Fig. S5.**
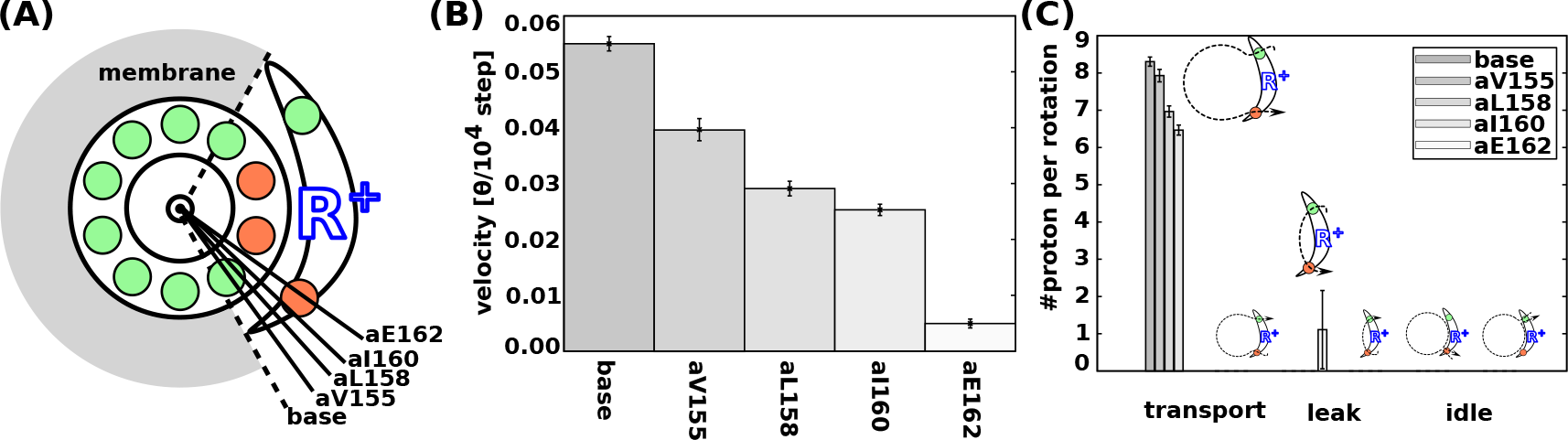
The effect of the lipid membrane border on the F_O_ rotation. (A) Schematic picture of the altered lipid membrane regions. The concrete angles are −69.4°, −59.1°, −52.2°, −48.4°, and −40.6° degree from the base to asub137, respectively. (B) The rotational velocity of each setup. The more the lipid membrane region intrudes a-subunit, the slower it rotates. (C) The number of protons moved per one rotation. The number of transported protons decreases as the membrane region expands. Additionally, proton leaks without contributing to the F_O_ rotation when the membrane region locates at the aE162 (asub137).

**Table S1.** The number of proton –transfer per rotation.

| # of proton per rotation | Transport |  | Idle |  |
| --- | --- | --- | --- | --- |
|  | IMS→matrix | matrix→IMS | IMS→IMS | matrix→matrix |
| Synthesis | 10.0±0.1 | 0 | 0 | 0 |
| Hydrolysis | 0 | -10.2±0.15 | 0.1±0.038 | 0 |

In the both setups have no leak proton so it was omitted.

**Table S2.**
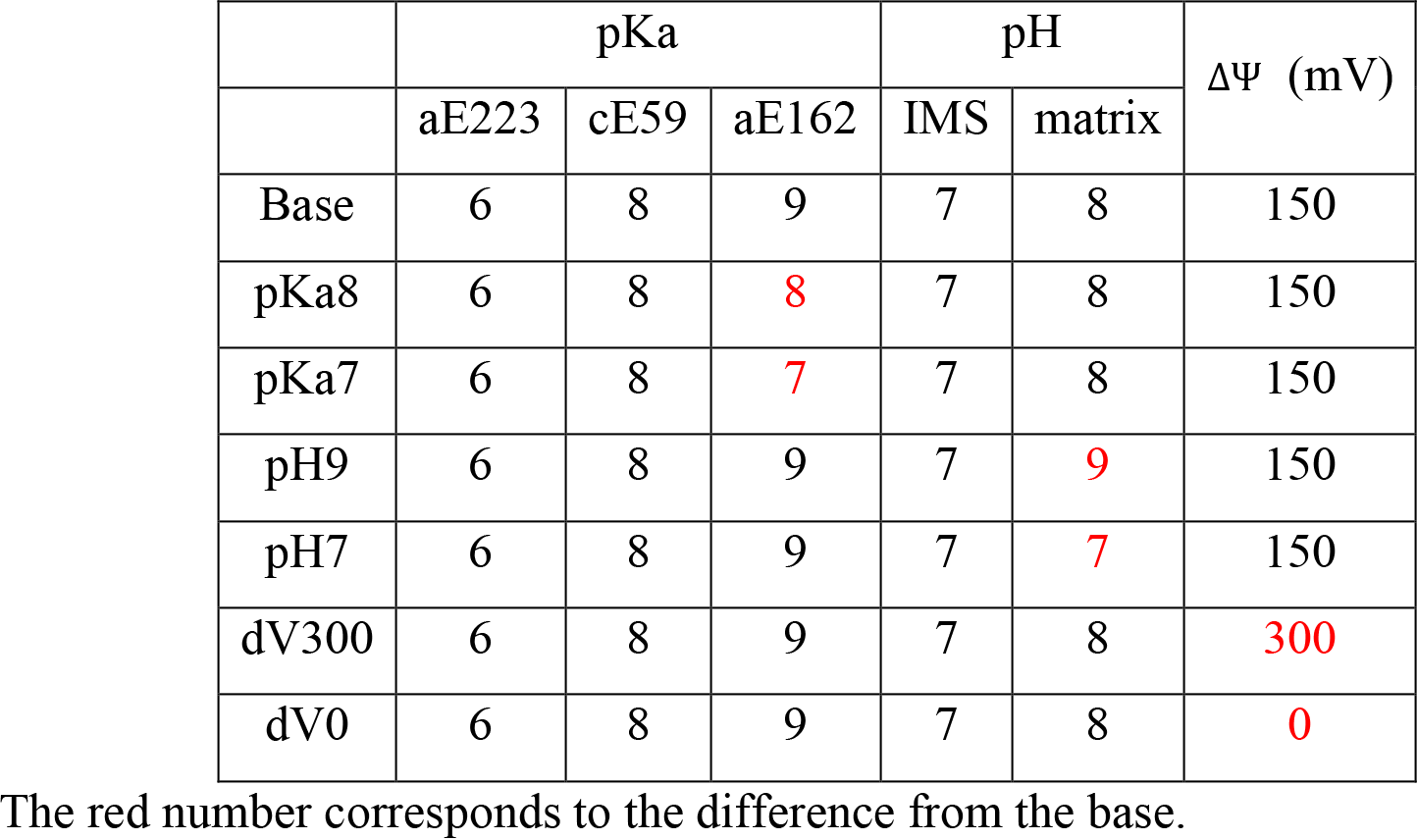
The list for Fig. 5.

| | pKa | | | pH | | $\Delta\Psi$ (mV) |
| --- | --- | --- | --- | --- | --- | --- |
|  | aE223 | cE59 | aE162 | IMS | matrix |  |
| Base | 6 | 8 | 9 | 7 | 8 | 150 |
| pKa8 | 6 | 8 | 8 | 7 | 8 | 150 |
| pKa7 | 6 | 8 | 7 | 7 | 8 | 150 |
| pH9 | 6 | 8 | 9 | 7 | 9 | 150 |
| pH7 | 6 | 8 | 9 | 7 | 7 | 150 |
| dV300 | 6 | 8 | 9 | 7 | 8 | 300 |
| dV0 | 6 | 8 | 9 | 7 | 8 | 0 |

The red number corresponds to the difference from the base.

**Movies S1 to S2.** The coloring method is same as Fig. 1

